# PRMT1-mediated methylation of the Large Drosha Complex regulates microRNA biogenesis

**DOI:** 10.1101/466813

**Authors:** Valeria Spadotto, Roberto Giambruno, Enrico Massignani, Marija Mihailovich, Francesca Patuzzo, Francesco Ghini, Francesco Nicassio, Tiziana Bonaldi

## Abstract

MicroRNA (miRNA) biogenesis is a tightly controlled multi-step process operated in the nucleus by the activity of the Large Drosha Complex (LDC). Through high resolution mass spectrometry (MS) analysis we discovered that the LDC is extensively methylated, with 82 distinct methylated sites associated to 16 out of 23 subunits of the LDC. The majority of these modifications occurs on arginine (R)- residues (61), leading to 86 methylation events, while 29 lysine (K)-methylation events occurs on 21 sites of the complex. Interestingly, both depletion and pharmacological inhibition of PRMT1 lead to a widespread alteration of the methylation state of the complex and induce global decrease of miRNA expression, as a consequence of the specific impairment of the pri-to-pre-miRNA processing step. In particular, we show that the reduced methylation of the ILF3 subunit of the complex is linked to its diminished binding to the target pri-miRNAs. Overall, our study uncovers a previously uncharacterized role of R-methylation in the regulation of the LDC activity in mammalian cells, thus affecting global miRNA levels.

## INTRODUCTION

MicroRNAs (miRNAs) are short non-coding RNA molecules that regulate gene expression at the post-transcriptional level [1–4]. They interact with target mRNAs by pairing with the corresponding miRNA-binding sites typically located in the 3’ untranslated regions (3’UTRs) and promote their translational repression and/or degradation [2]. MicroRNAs are preferentially transcribed by RNA Polymerase II into long primary transcripts, called pri-miRNAs, that possess the 7-methyl-guanosine cap at the 5’-end, the poly-A tail at the 3’-end and the stem-loop structures, where the mature miRNA sequences are embedded [5]. Genes encoding miRNAs are located in different genomic regions: intergenic miRNAs are transcribed as separated transcriptional units; while intragenic miRNAs are transcribed together with their “host” gene, the majority encoded within introns and a few deriving from exons [6]. Interestingly, miRNA loci located in close proximity are often co-transcribed as unique transcripts, giving rise to polycistronic units, composed of 2 to 19 individual miRNA hairpins [5,6].

In the nucleus, the Microprocessor complex, which comprises the type-III RNase Drosha and DGCR8, processes pri-miRNAs into shorter stem-loop molecules of 60-70 nucleotides, called precursor miRNAs (pre-miRNAs) [4,7]. DGCR8 binds to the pri-miRNA through its double strand RNA-binding domain and favors the correct positioning of the two RNase III domains of Drosha [8–10] which is a crucial step for the subsequent pri-miRNA cleavage and determination of the guide and passenger miRNA strands [11–14]. Pre-miRNAs are then exported in the cytoplasm by the exportin-5 (XPO5)- RAN-GTP complex and processed by the Dicer/Trbp complex into small RNA duplexes, about 22nt-long [15–18]. These duplexes are finally loaded into the RNA-Induced Silencing Complex (RISC), where the dsRNA is unwound, the passenger strand is removed and degraded, while the guide strand is retained and used for the recognition of the miRNA-binding site within the mRNA targets [19,20].

The tight control of microRNA biogenesis at multiple steps ensures the production of the correct levels of miRNA molecules that, in turn, fine-tune gene expression. Aberrant miRNA levels have been, in fact, observed in several pathologies, including cancer [21,22]. The main regulation of miRNA biogenesis is achieved through the modulation of the Microprocessor activity, which is rate-limiting for the whole process. The expression and activity of the Microprocessor is controlled in multiple ways: first, Drosha and DGCR8 protein levels are tightly regulated by a double-negative feedback loop, whereby DGCR8 stabilizes Drosha protein level, which, in turn, promotes the degradation of DGCR8 transcript by cleaving two hairpins located in its 5’UTR [23,24]. Second, although the Microprocessor alone can complete the pri-miRNAs cleavage reaction, there is evidence that various accessory proteins associate to it, and regulate its catalytic activity. So far, 21 co-factors have been described to interact with the Microprocessor, forming a bigger multi-protein complex called Large Drosha Complex (LDC). Accessory proteins comprise mainly RNA binding proteins (RBPs), such as the DEAD-box helicases DDX5 and DDX17, a number of heterogeneous ribonucleoproteins (hnRNPs), the FET proteins (FUS, EWSR, TAF15) and other factors [4,25,26]. They modulate the catalytic activity and define the substrate specificity of the Microprocessor, in various ways [4,25,27,28]: DDX5 and DDX17, for instance, are required for the recognition and processing of specific secondary structures within a subset of pri-miRNAs [27]; TARDBP both facilitates the binding and cleavage of specific pri-miRNAs and stabilizes Drosha levels by protecting it from the proteasome-dependent degradation [29,30]. ILF2 (also known as NF45) and the splicing isoform known as NF90 of ILF3 were initially considered negative regulators of miRNA biogenesis, being shown to sequester some pri-miRNAs (e.g. pri-let-7a and pri-miR-21) from the Microprocessor when overexpressed [31,32]. More recent experimental evidences based on gene knockdown experiment have, instead, demonstrated that basal ILF3 stabilizes specific pre- and mature miRNAs, thus exerting a positive regulation on the biogenesis of some miRNA [33].

The LDC can also be regulated by post-translational modifications (PTMs): phosphorylation and acetylation stabilize the levels of DGCR8 and Drosha, respectively [34,35], whereas Drosha phosphorylation by the MAPK p38 promotes its cytoplasmic export and degradation [36]. In 2013, our group discovered for the first time that several LDC subunits are methylated on arginine (R) residues [37], an observation that was further confirmed by a subsequent global methyl-proteomic study from another group [38]. However, besides the high frequency of this modification on the complex, its functional impact on the LDC-activity, and consequently miRNA biogenesis, remains elusive.

A family of 9 protein R-methyltransferases (PRMTs) can catalyze R-methylation in mammals. They are grouped in 3 classes based on the specific reaction that they can catalyze: type-I enzymes generate both mono-methylation and asymmetric di-methylation, whereby asymmetric di-methyl arginine (ADMA) present 2 methyl-groups added to the same terminal ω-guanidino nitrogen atom; type-II enzymes catalyze symmetric di-methylation at distinct terminal ω-guanidino nitrogen atoms of R (SDMA); type-III group comprises only PRMT7 that is capable of generating exclusively mono-methyl arginine (MMA) [39,40].

PRMT1 is the most active type-I PRMT, responsible of more than 85% of the annotated Rmethylations in the mammalian proteome [41,42], and it primarily modifies arginines located within the glycine- and arginine-rich sequences (RGG/RG) [43,44]. Besides the well-known target arginine 3 on histone H4 (H4R3me2a) [45,46], several non-histone proteins are substrates of PRMT1, such as the transcription factor RUNX1 [47]; the transcription elongation factor SPT5 [48]; some enzymes involved in the DNA damage response like MRE11, 53BP1 and BRCA1 [49–52]; and several RNA binding proteins, including also the LDC subunits ILF3, EWSR1, FUS and TAF15 [41,53–56]. It has been shown in various studies that PRMT1-dependent methylation can regulate proteins by affecting their subcellular localization, or interaction with both other proteins and nucleic acids, in particular RNAs [54,55,57–60]. Prompted by our initial evidence that LDC is hyper-methylated [37], we set to investigate the possible role of this modification in regulating both the composition and function of the complex and consequently, miRNA biosynthesis.

Overall, this study demonstrates for the first time that extensive R-methylation of the LDC is dependent on PRMT1 and impacts on the activity of the complex, thus regulating miRNA production and overall levels.

## RESULTS

### In-depth characterization of the LDC methyl-proteome

To obtain an accurate representation of all possible methylations occurring on the LDC, we combined the affinity-enrichment of the complex with heavy methyl (hm)SILAC-labeling and MS-proteomics. Methionine is an essential amino acid that, in the cell, is the precursor of Sadenosyl methionine (SAM), the sole donor of methyl-groups in enzymatic methylation reactions. In a hmSILAC context, cells are grown in media containing the heavy isotopic variant of methionine, ^13^CD_3_-methionine (Met4). Upon uptake, Met4 is intracellularly converted to ^13^CD_3_-SAM, which serves as donor of heavy methyl-groups for lysine-(K) and Rmethyltransferases so that substrate proteins become heavy methyl-labelled. When coupled to high resolution MS, this strategy allows distinguishing with high confidence *in vivo* methylation events on R and K from false positive identifications, such as chemical methylation, or amino acid substitutions that are isobaric to this modification [61]. In fact, in the MS spectra heavy methylated peptides result as isotopic peptide-pairs, where the heavy and the light peaks differ of unique mass-differences (deltamass, Δmass), on the basis of the number of methyl-groups added. The experimental workflow designed for the characterization of the LDC methyl-proteome is summarized in Fig. 1A. Based on the preferential localization of the LDC in the nucleus, the nuclear fraction of hmSILAC-labeled cells was used as input for the immuno-precipitation (IP) of the complex (Suppl. 1A). In order to maximize the LDC protein coverage, we performed 4 independent co-immunoprecipitations (co-IPs) using as bait 4 different LDC subunits: DDX5, DGCR8, Drosha, and FUS (Fig. 1B). In each IP, the specific bait was efficiently precipitated and the complex was enriched over background proteins (Fig. 1C, Suppl. 1B). All subunits were reproducibly identified in the immuno-precipitated material, with an average sequence coverage of more than 30% (Suppl. 1C). Upon protein separation, digestion and MS analysis of each IP fraction, MaxQuant output data was further analysed using the hmSEEKER pipeline, in-house developed to improve the identification of genuine methyl-sites from hmSILAC MS data. HmSEEKER searches for the isotopic peptide-pairs (doublets) among all detected MS1 peaks, allowing the identification of methyl-doublets even if only one of the two peak-counterparts is sequenced at the MS2 level (*Massignani E. et al., under revision*). Through this analytical pipeline, we identified 116 methylated peptides with high confidence (Suppl. Table 1), thus significantly extending our previous annotation [37]. The modified peptides correspond to 82 distinct R- or K-sites in total, associated to 16 out of 23 subunits of the LDC (Fig. 1E). Notably, the majority (75%) of these modifications occurs on R-residues, of which 40% are mono- and 60% di-methylated (Fig. 1F and 1G). In total 61 distinct R-sites are modified, of which 27 are exclusively di-methylated, 9 only mono-methylated and 25 both mono- and di-methylated (Fig. 1G). Individual IPs of four selected proteins of the LDC, followed by WB profiling using anti-pan-methyl-R antibodies confirmed that they exist as both mono- and asymmetrical di-methylated (Suppl. 1D). Interestingly, only 21 lysines within the LDC resulted methylated, of which 72% are mono-methylated, 14% di-methylated and 14% tri-methylated. All together 13 K-sites are exclusively mono-methylated; 4 K-sites can exist as both mono- and di-methylated (14%); and 4 K-sites are mono- and tri-methylated (14%) (Fig. 1G). Overall, the data indicate a prevalence of arginine modification within the complex.

**Figure 1.**
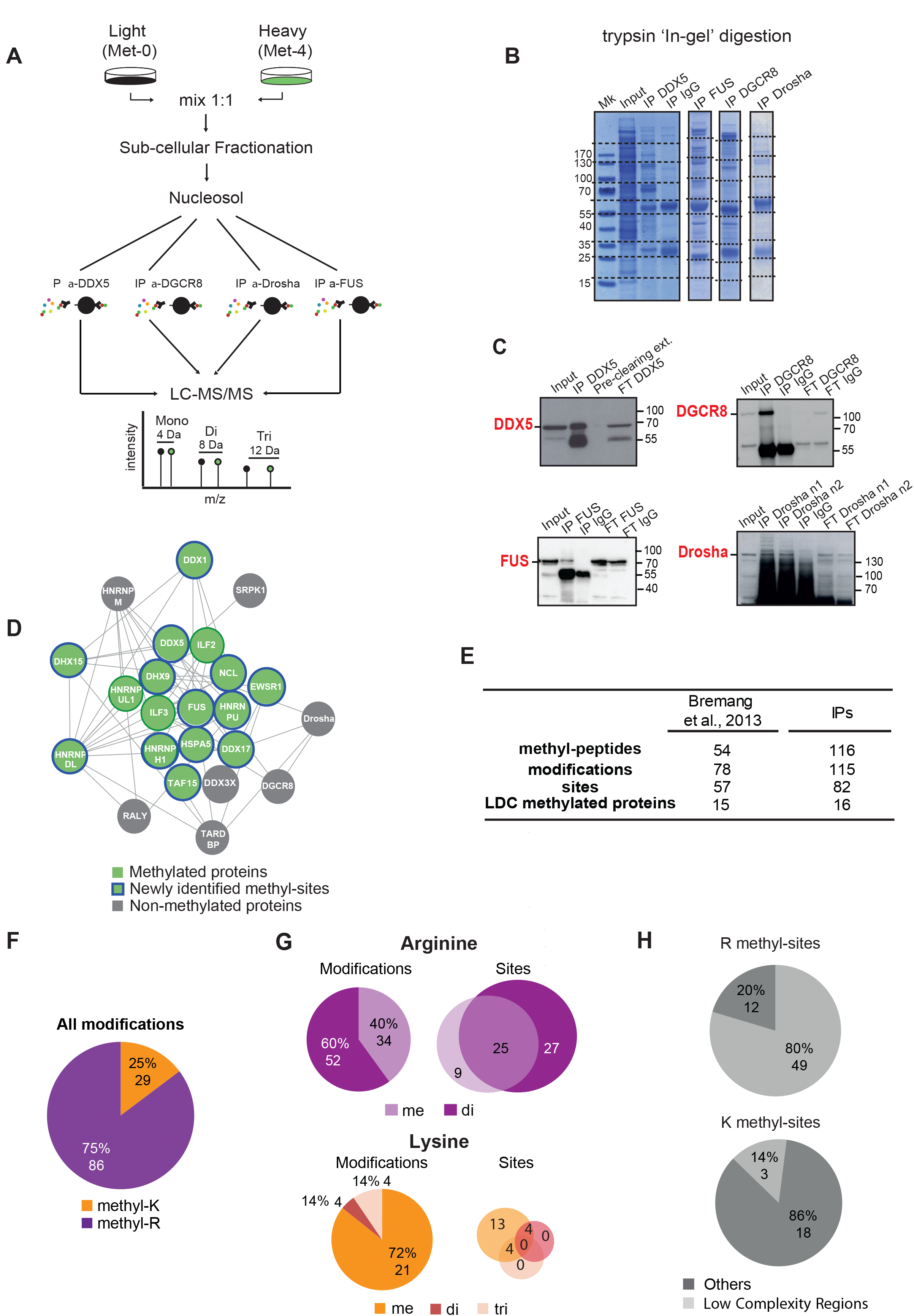
Characterization of the LDC methylation pattern by MS. **A)** Workflow of the hmSILAC/co-IP approach applied to characterize the LDC methyl-proteome. The methylation degree can be identified based on the mass difference between the light and heavy peptide (4 Da = mono-methylation, 8 Da = di-methylation, 12 Da = tri-methylation). **B)** Coomassie-stained gels of the immuno-precipitated material in-gel digested with trypsin and the analyzed by LC-MS/MS; dashed lines correspond to the individual gel slices processed and MS-analyzed. The acronym Mk stands for protein molecular marker. **C)** Validation of the efficiency of the IP of the 4 LDC proteins used as baits, by WB analysis. **D)** Graphical representation of the LDC complex using Cytoscape with methylated LDC subunits displayed in green, non-methylated LDC proteins in grey; blue circles indicate proteins bearing the newly annotated methyl-sites. **E)** Summary of the annotated LDC methyl-proteome, compared to one previously published [37]; **Methyl-peptides:** number of identified peptides harboring one or multiple methylation events. **Modifications:** total number of mono-, di- and tri-methylation events occurring on both K- and R-residues identified by hmSILAC/co-IP. **Sites:** number of methylated residues. **LDC methylated proteins:** number of LDC protein subunits found methylated by hmSILAC/co-IP. **F)** Summary of all arginine (R) and lysine (K) methylations on the LDC-complex. **G)** Upper panel left: summary of mono- and dimethyl-arginines identified by hmSILAC/protein IP. Upper panel right: Venn diagram of the R-sites identified as either mono- or di-methylated. Lower panel left: summary of the mono-, di- and trimethyl-lysines identified by hmSILAC/co-IP. Lower panel right: Venn diagram of the K-sites identified as mono-, di- and tri-methylated. **H)** Localization analysis of the R-methyl-sites (upper panel) and K-methyl-sites (lower panel) identified according to the SMART domain database [95].

Interestingly, 80% of R-methylated sites occur within Low Complexity (LC) regions (Fig. 1H), which have been often described to be involved in protein-protein and protein-RNA interactions [62–64]. Conversely, 86% of K-methylated sites are located outside of LC regions. This suggests that R-methylation may affect either the composition/stability of the complex or the binding of the LDC subunits with substrate miRNAs.

### PRMT1 down-regulation impairs global miRNA expression

To assess whether R-methylation regulates the LDC activity, we investigated the effect that modulating this PTM could exert on miRNA biogenesis. We focused on PRMT1 because it is the most active enzyme of the PRMT family, and generated HeLa cells knocked-down (KD) for this protein by RNA interference with two distinct short hairpin RNAs (shRNAs) inserted in lentiviral constructs. Reduction of the protein level was observed upon cell transduction with the two shRNAs (Fig. 2A). Up to 96h post-infection, the growth of PRMT1-depleted cells was similar to control cells, transduced with the empty vector (EV) (Suppl. 2A), whereas at later time points KD cells showed a progressive growth reduction until full arrest, in accordance with the reported embryonic lethality due to PRMT1 loss, in mice [42].

**Figure 2.**
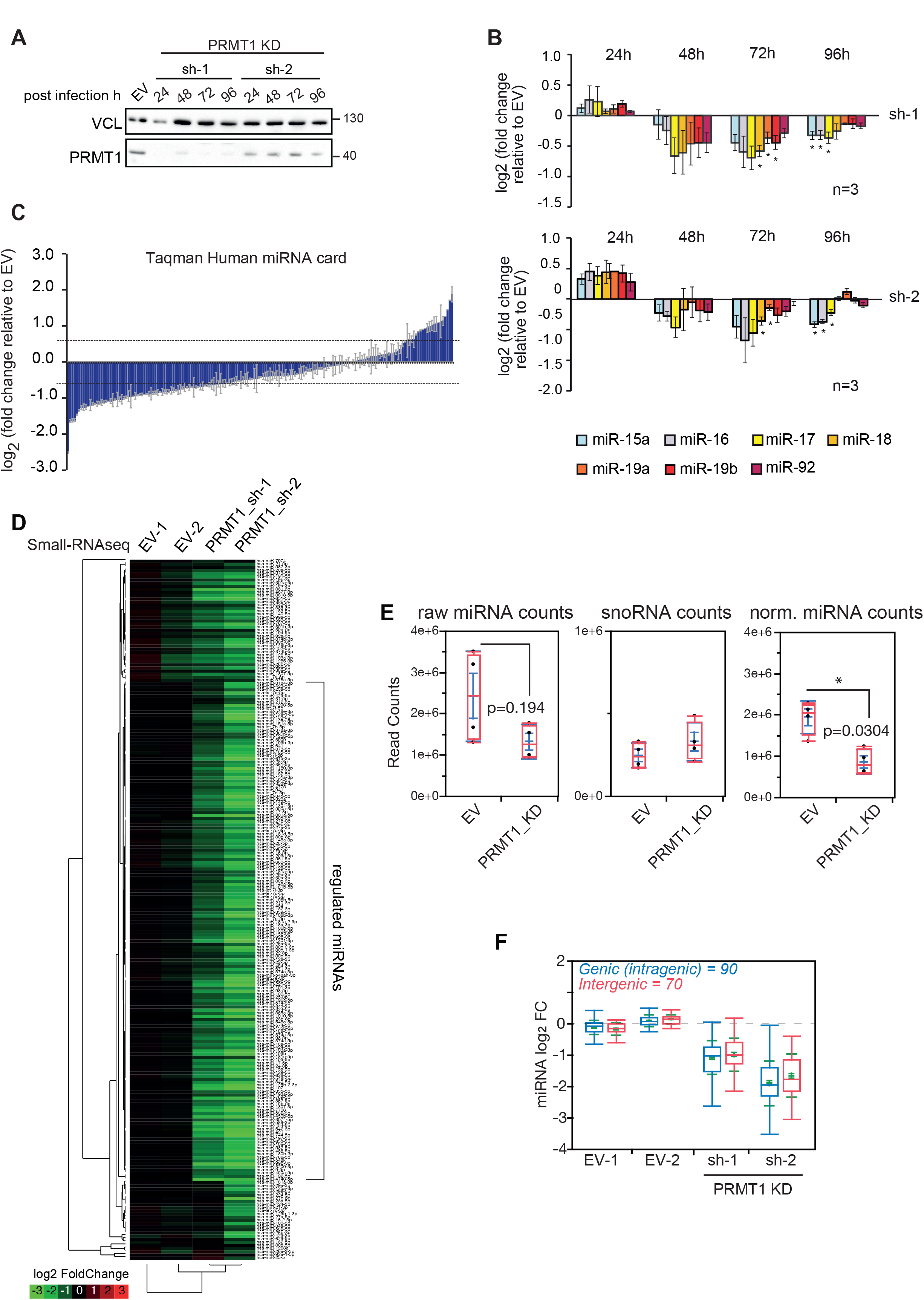
PRMT1 down-regulation impairs mature miRNA expression, globally. **A)** Time-course profiling of PRMT1 depletion by shRNA. The protein knock-down (KD) was obtained using two different shRNAs, sh-1 and sh-2 (see Material and Methods). VCL was used as loading control of protein level. **B)** Time-resolved expression profiling of mature miRNA levels of the miR-15a/16 and miR-17-92 clusters, upon PRMT1 depletion; data are presented as log_2_ fold enrichment of PRMT1 sh-1 (upper panel) and sh-2 (lower panel), over control (EV). Histograms represent mean ± SEM of 3 independent biological experiments (n=3). Statistical analysis was performed using non-parametric, 2-tailed T-Test. * = Values with a significant P-value < 0.01. **C)** Global expression analysis of miRNAs upon PRMT1 depletion by PRMT1 sh-1, performed with TaqMan Array Human miRNA Card. Data are displayed as log_2_ fold changes and are normalized on the geometric mean of a panel of housekeeping genes (mammU6, RNU44, RNU46, U6snRNA). Histograms represent mean ± SEM of miRNAs quantified in 2 technical replicates. Regulated miRNAs were considered significant when their fold change was greater/lower than 1.5. **D)** Heat map displaying log_2_ fold changes of miRNAs identified in control (EV) and PRMT1 KD achieved with the sh-1 and sh2 shRNA constructs. Data are normalized over the small nucleolar RNAs. The average of 2 technical replicates is shown for each sample. **E)** Box-plot of the distribution of miRNA and small nucleolar RNA (snoRNA) raw counts, and of miRNA normalized over snoRNA counts. Statistical analyses were performed using non-parametric Wilcoxon test. **F)** Box-plot shows the distributions of intragenic and intergenic miRNAs upon PRMT1 KD, obtained from miRiad [96]. Only guide miRNAs were considered for the analysis.

To assess the effect of PRMT1 depletion on LDC activity, we monitored the expression of two cancer-related miRNA clusters, miR-15a/16 and miR-17-92 [65,66], whose deregulation was mechanistically linked to the lack of some LDC subunits [67–69]. We observed a significant down-regulation of the mature miRNAs from both clusters, with this effect peaking at 72h post-infection (Fig. 2B). To achieve a more global view on miRNA biogenesis, we profiled miRNAs in PRMT1-depleted and control cells by both Small-RNAseq and qPCR using the Taqman human miRNA cards, 72h post-infection (Suppl. Table 2 and 3). Interestingly, we detected a pervasive miRNA down-regulation, with only a minor proportion of unchanged or up-regulated miRNAs (Fig. 2C, 2D). However, the effect on miRNAs was not mirrored by a change in small nucleolar RNAs (snoRNAs) (Fig. 2E), which indicates that PRMT1 depletion did not impact on other cellular ncRNAs. With both techniques, we confirmed the down-regulation of miR-15a/16 and miR-17-92 (Suppl. 2B).

To understand whether PRMT1 affects the biosynthesis of specific classes of miRNAs, we first focused on the analysis of intronic and intergenic miRNAs, whereby the former derive from the processing of introns and require spliceosomal components for their biogenesis, while the latter are transcribed as independent transcription units [70,71]. Observing no differential expression between the two miRNA types in PRMT1 deficient cells (Fig. 2F), we concluded that the reduction in miRNA processing does not depend on their genomic origin. Second, we analyzed the effect of PRMT1 on the positioning and orientation of Drosha cleavage, elaborating on the fact that miRNAs can originate from the 5‟ (5p) or 3‟ (3p) arm of the pri-miRNA hairpin depending on Drosha activity [14]. The observation that 5p and 3p miRNAs are equally regulated suggests that PRMT1 depletion does not affect directly the catalytic activity of Drosha towards one of the two arms (Supp. 2C).

### PRMT1 depletion impairs the processing of primary-to-precursor miRNAs

The primary function of the LDC is to cleave nuclear pri-miRNAs into shorter pre-miRNAs. Thus, we sought to investigate whether the global miRNA down-regulation observed upon PRMT1 depletion is caused by the specific impairment of this catalytic step. We designed quantitative real time PCR (qPCR) primers that allow distinguishing pri- from pre-miRNAs deriving from miR-15a/16 and miR-17-92 clusters and used them in combination to standard primers directed for the amplification of the mature forms (Fig. 3A). Quantitative PCR analysis indicated that pri-miR-15a/16 and pri-miR-17-92 levels were unchanged or even increased, upon PRMT1-depletion (Fig. 3B), whereas pre- and mature miRNAs were significantly reduced (Fig. 3C and 3D). To corroborate the evidence that PRMT1 KD affects specifically pri-to-pre-miRNA processing, we also profiled the pri- and pre-miRNAs from the miR-15a/16 and 17-92 clusters upon cell separation into chromatin and nucleosolic fractions, since pri-miRNAs are more associated to chromatin while pre-miRNA are released in the nucleosol prior to cytoplasmic export. By profiling pri-miRNAs in the chromatin fraction and pre-miRNAs in the nucleosol, we confirmed that pre-miRNAs are down-regulated upon PRMT1 KD while the corresponding pri-miRNAs are unchanged (Fig. 3E). Overall, these findings demonstrate a mechanistic link between PRMT1 expression levels and the reaction specifically mediated by the LDC.

**Figure 3.**
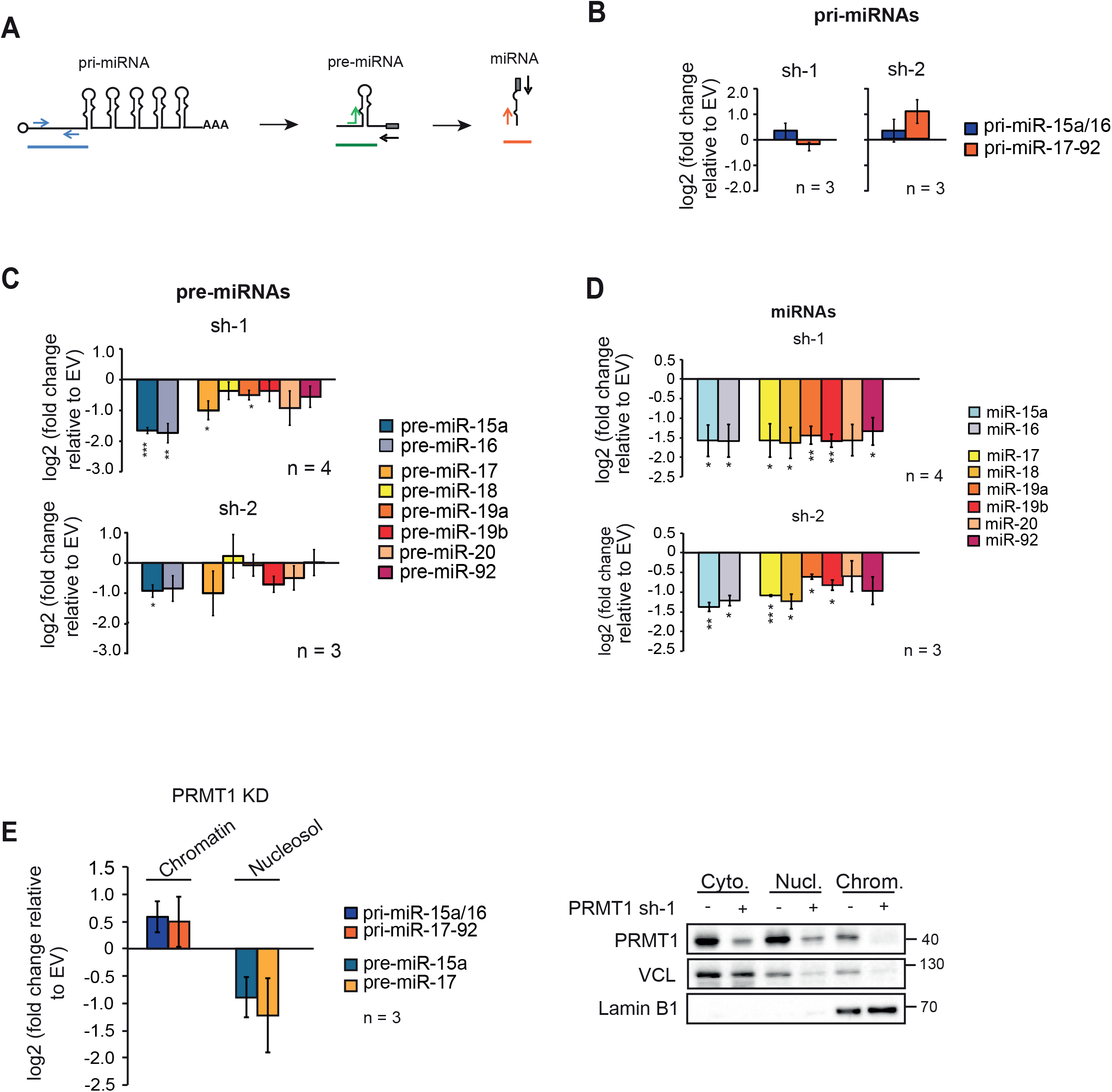
The processing of primary-to-precursor miRNAs is impaired upon PRMT1 depletion. **A)** Schematic representation of the qPCR primers designed for the selective detection of pri-, pre- and mature miRNAs. The arrows indicate the region that is amplified by qPCR for each miRNA isoform. The rectangle in grey indicates the binding region of the Qiagen universal primer which is colored in black and is used to amplify the same 3’end region of pre- and mature miRNAs according to the vendor protocol. **B)** qPCR profiling of pri-miRNAs of miR-15a/16 and miR-17-92 upon PRMT1 KD, achieved with the sh-1 and sh-2 shRNA constructs. Histograms represent mean ± SEM of the log_2_ fold change of PRMT1-depleted cells over EV control cells from 3 biological experiments (n=3) **C)** qPCR profiling of the pre-miRNAs from miR-15a/16 and miR-17-92, upon PRMT1 KD with sh-1 and sh-2. Histograms represent the mean ± SEM of the log_2_ fold change over EV control cells from the averages of 4 biological replicates for sh-1 (n= 4, upper panel) and 3 biological replicates for sh-2 (n= 3, lower panel). **D)** qPCR profiling of mature miRNAs derived from miR-15a/16 and miR-17-92 clusters upon PRMT1 KD with sh-1 and sh-2. Histograms represent the mean ± SEM of the log_2_ fold change of KD over EV control cells. The results are the average of 4 biological replicates for sh-1 (n=4, upper panel) and 3 biological replicates for sh-2 (n=3, lower panel). Statistical analysis was performed using non-parametric, 2-tailed T-Test. * = Values with p-value < 0.05. ** = values with p-value ≤0.01 and *** = values with p-value ≤0.001. **E)** Left panel: qPCR analysis of pri-miR-15a/16, pri-miR-17-92 and pre-miR-15a and pre-miR-17 in chromatin and nuclear fractions of PRMT1KD HeLa cells. The data are the average from 3 biological experiments (n=3) and expressed as log_2_ fold change over control. Right panel: WB validation of the cellular fractionation in cytosolic (Cyto.), nucleosolic (Nucl.) and chromatin (Chrom.) compartments, of both control and PRMT1 KD cells.

### Modulation of PRMT1 strongly affects the methylation state of the LDC

To gain insights in the mechanism of PRMT1 regulation of the LDC activity, we characterized the expression, composition and methylation state of the complex by combining SILAC-based proteomics with the modulation of PRMT1 expression levels (Fig. 4A). We first chose the optimal time-point to carry out these analyses, by WB-profiling mono- and asymmetric dimethylation of arginines in PRMT1 KD and OE cells compared to the respective controls (Suppl. 3A, Suppl. 3C). When PRMT1 was depleted, we observed a reduction in the level of ADMA, mirrored by an increase of MMA over time, with a peak between 72 and 96h after cell infection, in line with previous studies [72] (Suppl. 3A). On the contrary, when PRMT1 was overexpressed, we observed a positive correlation between the enzyme levels and that of ADMA and MMA starting from 48h over time, compared with EV control cells (Suppl. 3C). Both PRMT1 KD and OE cells showed the strongest changes in global protein methylation after 72h of modulation of PRMT1 expression. Interestingly, the timing of these changes corresponded to the one in which we observed the down-regulation of miRNAs (Fig. 2). Hence, 72h post infection was chosen as the time point at which carrying out the MS-proteomics experiment (Fig. 4A). HeLa cells were metabolically labeled with either the light or the heavy isotopic variants of lysine and arginine (Arg0 and Lys0, and Arg10 and Lys8 for the light (L) and heavy (H) channels, respectively). In the “Forward” setting, EV control cells were cultured in the L medium and PRMT1 KD/OE cells in the H medium, while in the “Reverse” experiment, the L and H channels were swapped. Once harvested and mixed in 1:1 ratio, cells were fractionated into nuclear and cytoplasmic extracts (Suppl. 3B and Suppl. 3D) and a minor fraction of the nuclear extract was directly subjected to MS analysis for protein profiling of LDC (Fig. 4A). The remaining part was used as input for the LDC immuno-enrichment, carrying out three co-IPs in parallel, using DDX5, DGCR8, and FUS as baits (Suppl. 3B). Each IP was performed in three biological replicates.

**Figure 4.**
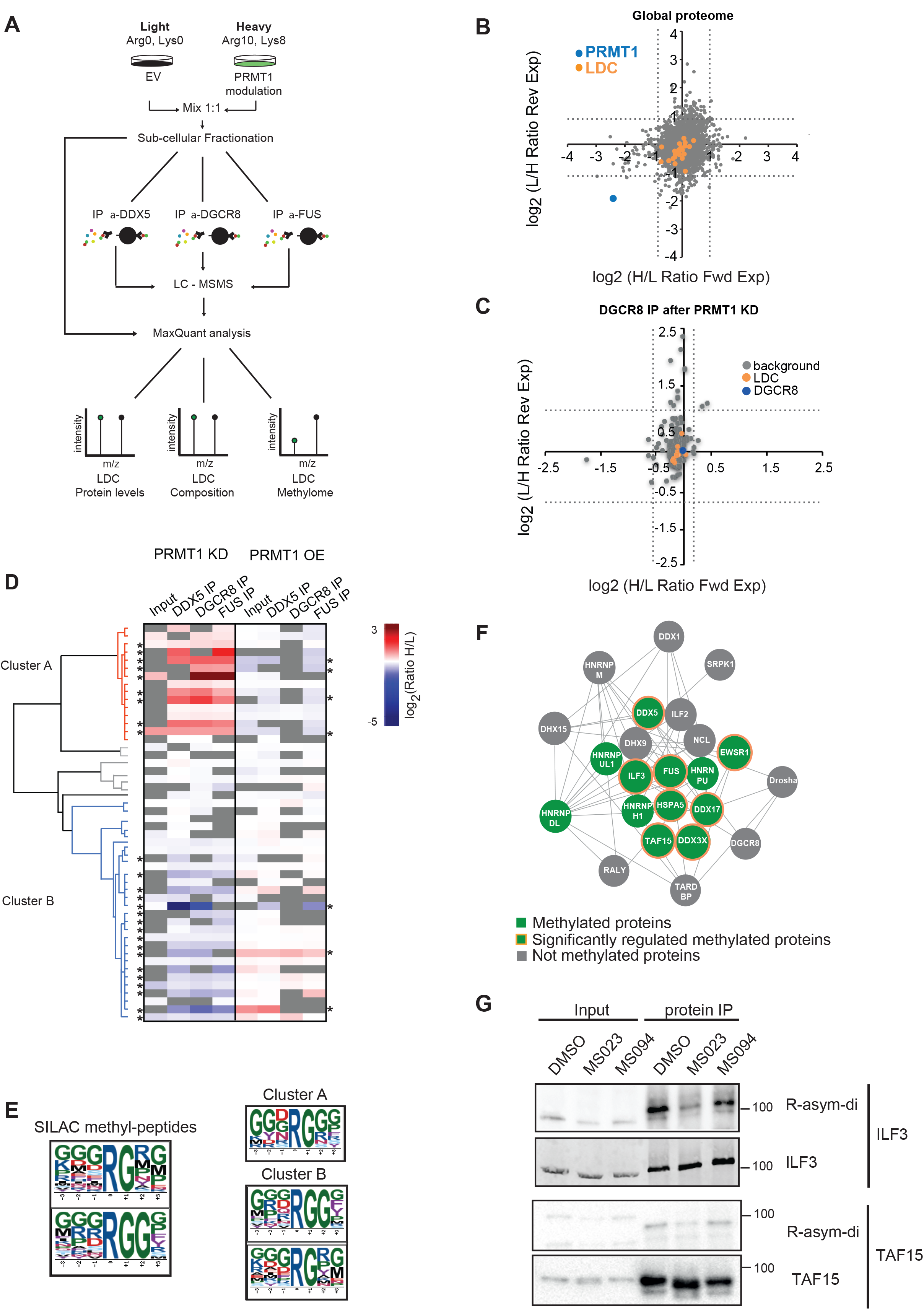
PRMT1 does not change LDC expression and composition but modulates LDC methylation. **A)** Workflow of the experimental setup designed for proteomics analysis of LDC expression and composition, upon PRMT1 modulation. Co-IP of the LDC using nuclear extracts from a 1:1 mix of SILAC-labelled HeLa S3 cells, depleted of- or overexpressing PRMT1 (KD, sh-1 and PRMT1 OE), or infected with the EV (control). Each immuno-enrichment of the complex with individual baits was carried out in 3 independent biological replicates (n=3), with SILAC channels swapping (Forward and Reverse experiments). **B)** LDC nuclear protein expression analysis: scatter-plot of all proteins identified and quantified in the SILAC nuclear inputs of HeLa cells, depleted or not of PRMT1. The scatterplot displays the log_2_ SILAC distribution of the quantified proteins in the Forward (Fwd) and Reverse (Rev) experiments (H/L ratio for the Fwd and L/H ratio for the Rev experiment. The dot corresponding to PRMT1 is highlighted in blue; LDC subunits are displayed in orange and all remaining proteins are in grey. Dashed lines indicate the μ±2Ϭ cut-off, calculated from the whole peptide SILAC ratio distribution in both Fwd and Rev experiments, that separate significant outliers form the unchanging population. **C)** LDC protein composition analysis: scatter plots of all proteins identified and quantified in the DGCR8 co-IPs, analyzed as described in B; the dot corresponding to DGCR8 is displayed in blue; all LDC subunits are indicated in orange, the remaining proteins are in grey. Dashed lines represent the μ±2Ϭ cut-off calculated on the protein ratio distribution, and define significantly changing form unchanging proteins. **D)** Heat map display of the unsupervised clustering analysis of methylated peptides identified and quantified in the SILAC/co-IP experiments upon PRMT1 KD and PRMT1 OE. Only peptides identified in at least 2 co-IP experiments of either the PRMT1 KD or PRMT1 OE were included in the analysis. Data are displayed as the average from 3 independent IPs for each bait and expressed as log_2_ ratio of PRMT1 KD/OE cells, compared to control cells. The heat map shows an unsupervised clustering of the data, generated using Pearson correlation and data average analysis. The 2 main clusters identified are indicated with differential color code (cluster A in red and cluster B in blue). *= Methylated peptides significantly regulated in the SILAC experiments upon a μ±1Ϭ cut-off calculated from the unmodified peptide distribution of each IP experiment. **E)** Motif enrichment analysis of the all SILAC-quantified methyl-peptides using ScanX software [97]. The following parameters were imposed: R central character; width= 7; occurrences= 10; significance= 0.000001 (right panel). The same analysis was performed also on the methyl-peptides identified in the cluster A and cluster B, separately (right panel). **F)** LDC complex representation with Cytoscape, of the proteins identified in the SILAC/co-IP experiments: in green are displayed proteins found methylated. Methylated proteins whose methylation state is significantly changing upon PRMT1 modulation are highlighted with orange circles. Non-methylated proteins are displayed in grey. **G)** Immuno-enrichment of ILF3 and TAF15 from HeLa cells treated with DMSO, the MS023 inhibitor and the MS094 control compound, used at 10μM concentration for 16 hours. WB profiling using anti-ADMA, anti-ILF3 and anti-TAF15 to assess the level of di-methylation and unmodified ILF3 and TAF15 protein levels, respectively.

**Figure 5.**
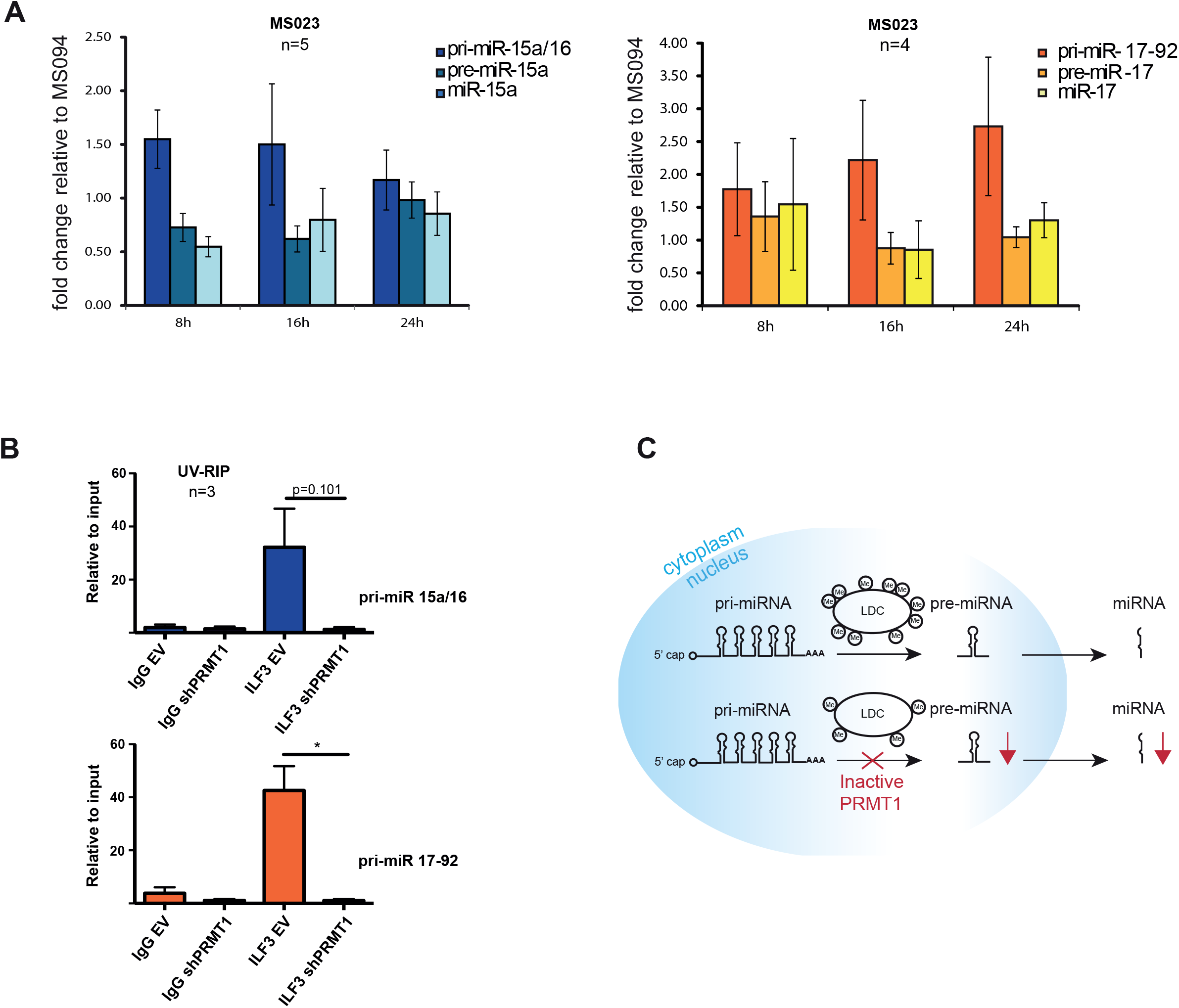
Pharmacological inhibition of PRMT1 inhibits the pri-to-pre miRNA processing step and affects ILF3 interaction with pri-miRNA targets. **A)** qPCR analysis of pri-, pre- and mature miRNAs from the miR-15a/16 and 17-92 clusters, upon treatment with MS023 and MS094 (10μM). Histograms represent the mean ± SEM of the fold change of MS023 over MS094 treated cells, calculated from 4 biological replicates (n=4). **B)** UV-RIP of ILF3 in HeLa cells expressing either the empty vector (EV) or the shRNA specific for PRMT1 (sh-1). IgG were used as mock control for the IP. The data shown are the average of 3 biological replicate experiments (n=3) and are represented as ratio over the input, thus indicating the fold enrichment. UV-RIP of ILF3 with pri-miR-15a/16 is shown in the upper bar-graph, while the with pri-miR-17-92 is in the bottom bar graph. Statistical analysis was performed using non-parametric, 2-tailed T-Test. * = Values with a significant P value < 0.1. **C)** Proposed model for the role of PRMT1 in miRNA biogenesis, through LDC methylation.

The SILAC ratios of the nuclear proteome upon PRMT1 KD did not indicate significant differences in the expression levels of the LDC subunits, compared to EV control (Fig. 4B), a result that was further confirmed by WB analysis of various LDC proteins carried out on the whole cell lysate of PRMT1 depleted cells (Suppl. 4A). In line with these results, no major changes were detected in the mRNA expression of most LDC subunits, except for HNRNPDL, SRPK1 and TAF15 whose transcript levels appeared slightly reduced upon PRMT1 knock-down (Suppl. 4B).

The stability of the LDC protein levels correlated with the fact that the overall composition of the complex, independently evaluated upon DGCR8, DDX5 and FUS co-IPs, remained substantially unaltered upon PRMT1 depletion (Fig. 4C and Suppl. 4C). In fact, the SILAC ratios of all subunits within each co-IP were similar to 1, indicating that the proteins were reproducibly co-immuno-precipitated within the complex using different baits. Considering that no major effects are observed at the level of LDC expression and composition after PRMT1 depletion, we hypothesized that the observed PRMT1-dependent impairment of miRNA biogenesis could be due to the alteration of the methylation state of the LDC.

We therefore analysed the methyl-proteome of the LDC upon PRMT1 KD and OE by quantitatively profiling the methyl-peptides by SILAC, in both the nuclear extract and the immuno-enriched complex. For the analysis of the LDC methylation state, we considered only methyl-peptides which were reproducibly identified in at least 2 out of 3 co-IPs (Fig. 4D) and identified 50 methyl-peptides displaying differences between PRMT1-modulated and control cells, of which 26 resulted statistically significant (Fig. 4D and Suppl. Table 4). The number of regulated methyl-peptides is higher in the KD than in the OE cells, probably due to the fact that the basal activity of the endogenous PRMT1 is already saturating the methylation at some sites, which are therefore not free for further modification by the exogenous enzyme. Moreover, the regulated methyl-sites are high-quality identifications, since 80% was orthogonally validated by hmSILAC (Suppl. 5A, upper part). Interestingly, the number of hmSILAC-validated methyl-peptides increases when only the significantly regulated peptides are considered (92%) (Suppl. 5A, lower part), which confirms on the one hand that the dynamic change of methyl-peptides is a good indicator of *in vivo* methylation and, on the other hand, that our high-confidence methylproteome is not yet saturated.

Unsupervised clustering analysis of the PRMT1-dependent LDC methyl-proteome showed the existence of 2 methyl-peptide clusters: cluster A collects peptides that are hyper-methylated upon PRMT1 depletion and hypo-methylated in PRMT1 overexpressing cells; cluster B includes peptides hypo-methylated in PRMT1 KD cells and whose methylation increased when PRMT1 is upregulated (Fig. 4D and Suppl. Table 5). Motif enrichment analysis of all significantly regulated methyl-peptides displayed a strong enrichment of both the RGG and RG motifs, where the former is more specific for PRMT1 and the latter is a more general target sequence for the whole PRMT family [40,73] (Fig. 4E, left panel). However, when we carried out the motif analysis separately on the peptides from the two clusters, cluster B displayed the enrichment of both motifs, whereas cluster A showed the enrichment of the sole RG motif (Fig. 4E, right panel and Suppl. Table 5). The observation that cluster B displays both the expected trend of variation in dependence of PRMT1 and the RGG consensus motif, more specific for PRMT1, suggests that the corresponding methyl-sites are more likely genuine PRMT1 targets. On the contrary, the peptides in cluster A - hyper-methylated upon PRMT1 depletion and enriched for the more generic consensus sequence- may be substrates of other PRMTs when the major enzyme of the family is lacking. In cluster B, we found DDX17, DDX3X, DDX5 and HNRNPH1 proteins, hereby identified as new PRMT1 targets (Suppl. 5B). Overall by quantitative proteomics we identified and characterized the exact methyl-sites of 12 subunits of LDC, 8 of which are also significantly modulated in dependence of PRMT1 (Fig. 4F).

To confirm the MS-data, we used the recently described type-I PRMTs inhibitor MS023 and its homologous inactive counterpart MS094 [74]. Treatment with MS023 led to a reduction of ADMA and a parallel increase of MMA, while MS094 did not induce methyl-proteome changes (Suppl. 5C). When we carried out the protein-IP of ILF3 and TAF15 followed by WB profiling with pan-methyl antibodies we observed that their basal di-methylation level was reduced upon MS023, while remained unaltered MS094, in agreement with the proteomic data (Fig. 4G). Instead, with the same approach we confirmed that DDX17 is exclusively R-mono-methylated and this modification is unaffected by MS023 (Suppl. 5D).

The results collected showing that LDC methylation is significantly affected by the modulation of PRMT1 expression or activity hints towards a key role of this modification in regulating the function of the complex and, consequently, miRNA biosynthesis.

### Inhibition of PRMT1 catalytic activity impairs miRNA biogenesis by reducing the interaction of LDC components with pri-miRNAs

To confirm that PRMT1 enzymatic activity, rather than its expression, is crucial for the correct processing of miRNAs, we assessed the levels of the intermediate products of miRNA biogenesis upon HeLa cells treatment with MS023 and MS094. A time-course inhibition experiment showed that MS023 exerts its effect on global ADMA/MMA levels already 8 hours upon drug administration (Suppl. 5E). We thus profiled the expression of primary, precursor and mature miRNAs of miR-15a/16 and miR-17-92 clusters in cells treated with the drugs at different time points and observed that the levels of pre- and mature miRNAs were reduced while the pri-miRNAs were unchanged at 8 and16 hours post inhibition (Fig. 6A). This confirms that the specific impairment of the pri-to-pre-miRNA processing step is due to the inhibition of the LDC catalytic activity.

Most of the LDC subunits identified as PRMT1 substrates by proteomics are RBPs. Moreover, their methylations occur within the RGG-rich sequences comprised in low complexity regions known to be involved in protein-RNA interactions [75]. Thus, we hypothesized that PRMT1-dependent R-methylation could modulate the interaction of some of the LDC subunits with their pri-miRNA targets. We focused on ILF3 because this protein emerged as a genuine PRMT1 target by MS, with a single di-methylated site on R609 which was significantly down-regulated upon PRMT1 KD and orthogonally-validated by hmSILAC (Suppl. 6A). We set up an experiment of UV-crosslinking followed by RNA-immunoprecipitation (UV-RIP) using ILF3 as bait, in control and PRMT1 KD cells and assessed the binding of ILF3 to the pri-miRNA-15a/16 and pri-miRNA-17-92. The observation that the binding of ILF3 to both pri-miRNAs is strongly reduced upon PRMT1 depletion (Fig. 6B) directly links PRMT1-mediated methylation of this LDC subunit with its interaction to the pri-miRNA substrates. These data allow formulating a model whereby PRMT1 regulates miRNA biogenesis through extensive R-methylation of the Large Drosha Complex, and in which its depletion or inactivation generate aberrant miRNAs expression (Fig. 6C).

## DISCUSSION

In this study, we took advantage of quantitative MS-based proteomics to investigate the extent, dynamicity and functional impact of R-methylation within the Large Drosha Complex. We initially combined MS-analysis with the affinity-enrichment of the complex to carry out a thorough characterization of its *in vivo* methylations. MS-analysis of protein methylation is particularly challenging both because this modification is isobaric to various amino acid substitutions and since chemical methylations introduced during sample preparation can be mis-assigned to the *in vivo* enzymatic modification. Thus, it has been demonstrated that label-free approaches for global analysis of methylation by MS can lead to high false discovery rate (FDR) in the absence of orthogonal validation strategies [76,77]. Yet, the generation of robust and reliable methyl-sites dataset is crucial for biological and functional follow up studies. We took advantage of the hmSILAC labeling-currently considered the golden standard strategy to reduce the FDRs in global MS-based identification of methyl-sites [77]- to generate the largest high-quality LDC methyl-proteome annotated so far.

Non-histone protein methylation, likewise other PTMs, is sub-stoichiometric and thus its analysis requires enrichment steps prior to MS. This is achieved by affinity purification strategies using pan-methyl antibodies, recently developed for global protein methylation analysis [37,76,78,79]. The methyl-proteomes annotated so far using this strategy indicate MMA as the most abundant methylation type present globally. While this may truly reflect the larger extent of this methylation degree over the others, it is still difficult to rule out whether instead it is consequence of the better performance of the anti-pan-MMA antibody [79]. As a matter of fact, the quantification of the different types of methylation carried out in MEFs in the absence of affinity-enrichment steps indicated ADMA as the predominant modification [72]. In line with this evidence, we found that 60% of the methyl-arginines within the LDC are di-methylated. This suggests that pan-methyl-antibodies may introduce biases in methyl-proteomics analyses and that the protein co-IP in combination with hmSILAC-MS may lead to more realistic pictures of the extent and composition of sub-methyl-proteomes.

Upon PRMT1 modulation, we observed significant changes in the methylation state of various LDC subunits that were thus identified for the first time as PRMT1 substrates, when their methylation state positively correlated with the enzyme level and occurred within the PRMT1-specific consensus motif RGG. Interestingly, our quantitative proteomic analysis also showed some peptides whose methylation was unaffected by changes of PRMT1 expression: they may be either targets of other PRMTs or constitutively methylated peptides possibly required for the correct assembly or functionality of the complex. Another subset of peptides was up-regulated in the absence of PRMT1 and enriched in the RG-motif: these may be substrates of other PRMTs, in line with the scavenging effect previously described upon PRMT1 ablation [72].

When analyzing the LDC methyl-proteome, we noticed that none of the identified methylation events occur in the nuclear localization signals (NLSs) of EWSR1, FUS and TAF15 that were previously reported to modulate their cytoplasm-to-nucleus translocation [54,80,81]. In line with this, HeLa treatment with MS023 and MS094 did not affect the subcellular localization of these proteins, confirming that the methyl-sites involved in the LDC function are different from the ones defining the protein localization (Suppl. 6B).

Conversely, we discovered that 80% of R-methylations occur on LC regions enriched in RG-sequences and typically involved in the interaction with RNA. According to the fact that the majority of the LDC proteins are RNA-binding proteins, we linked the wide-spread down-regulation of miRNAs upon PRMT1 ablation to a possible role of R-methylation in regulating the RNA-binding properties of the LDC subunits. Our hypothesis was confirmed by the observed reduced binding of ILF3 to pri-miR-15a/16 and pri-miRNA-17-92 upon PRMT1 depletion. It is likely that this molecular effect of protein-R-methylation may be more general, affecting the RNA-binding ability of other subunits of the complex that possess RG-rich sequences, as well as other RBPs in general. Even if our findings on ILF3 suggest that R-methylation promotes protein-RNA interactions, we cannot exclude that this modification may on the contrary exert an opposit effect on other subunits of the complex, as described for instance for hnRNPA1 [82]. In fact, although methylation does not change the net charge of the arginine guanidine group, it enhances its hydrophobicity, thus enabling two possible opposite effects: on the one hand, it could interfere with the hydrogen bonds established between arginine residues and the negatively charged phosphate backbone of the RNA, thus introducing steric hindrance for protein-RNA binding [83]; on the other hand, methylation could facilitate the stacking of methylated arginines within the bases of RNA, hereby promoting protein-RNA interactions [43].

The emerging role of PRMT1-dependent R-methylation on miRNA biogenesis is similar to that reported for other PTMs decorating the LDC, i.e. phosphorylation and acetylation, which regulate the intracellular levels of miRNAs in response to intra- and external-stimuli [34–36]. Interestingly, PRMTs are de-regulated in several types of cancer [39] and in particular PRMT1 overexpression was shown to positively correlate with tumor progression [84,85]. Also, a recent study has showed that the overexpression of type-I PRMTs, including PRMT1, promotes mammary gland tumorigenesis in mice [86]. Similarly, microRNA expression is often altered in cancer, where these molecules can display either oncogenic or tumour-suppressive functions [87–89]. In addition, proteins involved in miRNA biogenesis, such as Drosha and DGCR8, have also been found mutated and de-regulated in tumours [22], confirming the connection between aberrant LDC function, altered miRNA levels and cancer progression [20].

In this scenario, the fact that the miRNA biogenesis depends on R-methylation and can be modulated by PRMT inhibitors hints towards the possibility of using the pharmacological inhibition of PRMTs to control miRNA levels, offering novel therapeutic perspective for cancer treatment.

## MATERIALS AND METHODS

### Cell culture, Heavy methyl SILAC and SILAC labeling of cells

HeLa and HEK293 cells were grown in DMEM (Lonza, LA-0009E) supplemented with 10% FBS (Euroclone, ECSO182L), 1% glutamine (Lonza, BE17605E) and 100 U/mL Penicillin and Streptomycin (Euroclone, ECB3001D).

For Heavy methyl SILAC labeling, HeLa S3 cells were cultured in “Light” and “Heavy” SILAC media (PAA, custom) depleted of lysine, arginine and methionine and supplemented with L-arginine (Sigma Aldrich, A6969) L-lysine (Sigma Aldrich, L8662), and either L-[^13^CD3]-methionine (Met-4, heavy, Sigma Aldrich, 299154) or L-[^12^CH3]-methionine (Met-0, light, Sigma Aldrich M5308), as previously described (Ong SE et al., 2004).

For standard SILAC labeling, HeLa cells were grown in “Light” and “Heavy” SILAC DMEM (Thermo Fisher Scientific 88420) supplemented with either L-arginine and L-lysine, or their heavy isotope-counterparts L-arginine-^13^C_6_, ^15^N_4_ hydrochloride (Arg10, Sigma 608033) and L-lysine-^13^C_6_, ^15^N_2_ hydrochloride (Lys 8, Sigma 608041) (Ong SE et al., 2002).

All media were supplemented with 10% dialyzed FBS (26400-044 Gibco, Life Technology), 1% glutamine, 100 U/mL Penicillin and 100 mg/ml Streptomycin and 10 mM HEPES pH 7.5.

### Antibodies and chemical compounds

The following antibodies were used for immuno-precipitation and Western Blot experiments, according to the manufacturer’s instruction: DGCR8 (Abcam ab90579), EWSR1 (Abcam ab54708), FUS (Bethyl A300-293A), DDX5 (Abcam ab126730), Drosha (Santa Cruz sc-33778), VCL (Millipore 06-866), PRMT1 (Abcam ab73246), PRMT4 (Bethyl A300-421A), PRMT5 (Abcam ab109451), PRMT6 (Abcam ab47244), PRMT7 (sc-98882), ASYM24 (Millipore), SYM10 (Millipore), MeR4-100 (Cell Signaling Technology 8015), LAMIN A/C (sc-6215), Lamin B (Abcam ab16048), GAPDH (Abcam ab9484), H3 (Abcam ab1791), H4 (Abcam ab7311), H4R3me2a (Active Motif 39705), TAF15 (Bethyl Laboratories A300-308A), ILF3 (Bethyl A303-615A), ILF2 (sc-271718), DDX17 (sc-130650), DDX3 (LS-C64573).

MS023 and MS094 compounds were kindly provided by the SGC Toronto - Structural Genomic Consortium (http://www.thesgc.org/scientists/groups/toronto). Compounds were dissolved in DMSO and used at a final concentration of 10μM for the indicated timings.

### Cell transduction with lentiviral vectors

PRMT1 knock-down (KD) cells were generated with a second-generation pLKO lentiviral vectors, in which specific sh-RNA targeting PRMT1 were cloned. The pLKO-Empty vector was used as control.

To overexpress PRMT1, the cDNA of the v2 isoform was amplified starting from HeLa cDNA, using the following primers:

FW 5’–GGGGATCCCCCGGGCTGCAGATGGCGGCAGCCGAGGCCGCGAACTGCA– 3’ REV 5’–GAGGTTGATTGTCGACTCAGCGCATCCGGTAGTCGGTGG– 3’

The cDNA of PRMT1 was cloned into the pCCLsin.CPPT.PGK.GFP.WPRE lentiviral vector after plasmid linearization with PstI and SalI using the in-fusion HD EcoDry cloning plus, according to the manufacturer’s instructions (Clontech Laboratories, Inc., CA). The GFP cassette was removed from the vector upon digestion with XhoI.

### Cell lysis and sub-cellular fractionation

For preparation of whole cell extracts, cell pellets were lysed in 3 volumes of SDS Lysis Buffer (0.1 M Tris-HCl pH 7.5, 4% SDS), previously warmed to 95°C. Lysates were then sonicated, centrifuged 15 min at 13000 rpm to precipitate cell debris and then loaded on SDS-Page for subsequent protein separation.

For the preparation of cellular sub-fractions, cells were harvested, washed once with PBS and resuspended in 2 volumes of Lysis Buffer A (10 mM Hepes-KOH pH 7.9, 1.5 mM MgCl2, 10 mM KCl, 0.2% NP-40, 1X Roche Protease Inhibitors, 1U/μL NEB RNAse Inhibitors). After 20 strokes with a dounce homogenizer, cells were centrifuged 15 min at 3750 rpm. The supernatant (representing the cytoplasmic extract) was collected and the pellet (corresponding to crude nuclei) was washed twice with PBS and re-suspended in 2 volumes of Buffer C (420 mM NaCl, 0.1% NP40, 0.5 mM DTT, 20 mM Hepes-KOH pH 7.9, 2 mM MgCl2, 0.2 mM EDTA and 20% glycerol, 1X Roche Protease Inhibitors, 1U/μl NEB RNAse Inhibitors). The suspension was rocked 1h at 4°C and then ultracentrifuged at 33000 rpm for 1h.

For subsequent small RNA analysis from the same fractions, the chromatin pellet was washed with Buffer C, resuspended in Ripa Buffer (10 mM Tris-HCl pH 8, 150 mM NaCl, 0.1% SDS, 1% Triton, 1 mM EDTA, 0.1% Na Deoxycholate) and centrifuged 5 min at 13000 rpm at 4°C.

### Protein co-immunoprecipitation and Western Blot analysis

Experiments of protein co-IP were performed starting from 1-2 mg of whole cell extract in Ripa Buffer supplemented with fresh PMSF and 1x Roche protease inhibitors. When using nuclear extracts as input, 1-2 mg nuclear extracts were diluted with Ripa Buffer lacking NaCl, in order to decrease NaCl concentration to 150mM. The extracts were pre-cleared 3 times with 50 μL of Dynabeads protein G (Thermo Fisher Scientific) and then the specific antibody was added and incubated overnight on a rotating wheel at 4°C. The following day, 50 μL of Dynabeads protein G pre-equilibrated in PBS supplemented with 0.5% BSA, were added to the extract and incubated for 2 h on a rotating wheel at 4°C. Beads were then washed 3 times with Ripa Buffer and then bound material was eluted by incubation with LSD Sample Buffer (NuPAGE) supplemented with 100 mM DTT, at 95°C for 5 minutes. Samples were then loaded on SDS-PAGE for subsequent WB analysis.

### In-gel digestion of immunoprecipitated proteins

In gel digestion of gel-separated proteins with Trypsin, prior to MS analysis, was carried out as previously described [90]. After digestion and extraction from the gel pieces, the digested peptides were desalted and concentrated by reversed-phase chromatography onto micro-column C18 Stage Tips [91]. Peptides were then eluted from the tips with high organic solvent (80% ACN, 0.5% acetic acid), lyophilized, re-suspended in 1% TFA and subjected to LC-MS/MS analysis.

### Liquid Chromatography and Tandem Mass Spectrometry (LC-MS/MS)

Peptide mixtures were analyzed by online nano-flow liquid chromatography tandem mass spectrometry using an EASY-nLC™ 1000 (Thermo Fisher Scientific) connected to a quadrupole/Orbitrap mass spectrometer (Q Exactive, Thermo Fisher Scientific) through a nanoelectrospray ion source. The nano-LC system was operated in one column set up with a 25-cm analytical column (75 μm inner diameter, 350 μm outer diameter), packed with C18 reversed-phase resin (ReproSil, Pur C18AQ 1.9 μm, Dr. Maisch, Germany) configuration. Solvent A was 0.1% formic acid (FA) and 5% ACN in ddH_2_O and solvent B was 80% ACN with 0.1% FA. Peptides were injected at a flow rate of 500 nL/min and separated with a gradient of 5– 40% solvent B over 90 min, followed by a gradient of 40–60% for 10 min and 60–80% over 5 min at a flow rate of 250 nL/min. The Q-Exactive was operated in the data-dependent mode (DDA). HCD-fragmentation method when acquiring MS/MS spectra consisted of an Orbitrap full MS scan followed by up to 10 MS/MS experiments (Top10) on the most abundant ions detected in the full MS scan. Mass spectrometer conditions for all experiments were as follows: full MS (AGC 3e^6^; resolution 70,000; m/z range 300-1650; maximum ion time 20 ms); MS/MS (AGC 17,500; maximum ion time 50 ms; isolation width 1.8 Da with a dynamic exclusion time of 20 sec). Singly charged ions and ions for which no charge state could be determined were excluded from the selection. Normalized collision energy was set to 28%; spray voltage was 2.2 kV; no sheath and auxiliary gas flow; heated capillary temperature was 275 °C; S-lens RF level of 60%.

### Assigning hmSILAC/SILAC peptide sequences using MaxQuant and data analysis

Acquired Raw data were analyzed with the integrated MaxQuant software v.1.5.5.1 and v.1.6.0.16, using the Andromeda search engine [92,93]. The January 2016 version (UniProt Release 2016_01) of the Uniprot sequence was used for peptide identification. Enzyme specificity was set to Trypsin/P, meaning that trypsin cleavage occurs also in the presence of proline, after lysine or arginine residues. In MaxQuant, the estimated false discovery rate (FDR) of all peptide identifications was set to a maximum of 1%. The main search was performed with a mass tolerance of 6 ppm. A maximum of 3 missed cleavages were permitted, and the minimum peptide length was fixed at 7 amino acids. Carbamidomethylation of cysteine was set as a fixed modification.

### hmSILAC peptide assignment and data analysis

To assign hmSILAC peptide sequences, we defined new modifications in MaxQuant (v1.5.5.1 and v.1.6.0.16) with the mass increment and residue specificities corresponding to the heavy versions of mono-methylated K/R, di-methylated K/R, and tri-methylated K. Additionally, we defined new variable modifications for heavy methionine (Met4) and oxidized heavy methionine (Met4ox). To reduce the complexity of the search, and given the computational resources available, each experimental set of raw data was analysed three times using three distinct sets of variable modifications, namely: (1) Met4, Met4ox, oxidation, mono-methyl-K/R, mono-methyl4-K/R; (2) Met4, Met4ox, oxidation, di-methyl-K/R, di-methyl4-K/R; (3) Met4, Met4ox, oxidation, tri-methyl-K, tri-methyl4-K. Identification of high-confidence methylated sites was carried out with hmSEEKER, an in-house developed Perl pipeline that processes MaxQuant output files to find doublets of heavy and light hmSILAC peptides, by integrating the information contained in msms and allPeptides output files (*Massignani E. et al., manuscript under review*). HmSEEKER enables the retrieval of heavy/light methyl-peptide pairs whereby one of the two counterparts are not MS/MS identified. HmSEEKER performs the following steps: methyl-peptides identified in the evidence file are first filtered to remove: (1) all contaminants and decoy peptides; (2) all peptides with single charge and (3) all peptides bearing simultaneous heavy and light modifications. Then each peptide is associated to its corresponding MS1 peak in the allPeptides file. Finally, the H or L counterpart of each peak is searched among other peaks detected in the same raw data file. Because the pair is searched in msmsScans, hmSEEKER can find doublets even when one of the two counterparts has not been MS/MS sequenced, thus not appearing in the msms file. Assuming that the H and L counterpart must coelute, undergo the same ionization process and differ for a specific mass, we considered true positives the heavy/light peptide pairs that satisfied the following criteria: same charge, retention time difference between the two peptides <2 min and difference between observed and expected mass shift <7 ppm. In addition, hmSEEKER was used to automatically filter out all methyl-peptides carrying a modification with a localization probability <0.75. No Andromeda Score filtering was applied.

### Data analysis of identified SILAC peptides

SILAC peptide and protein quantification was performed automatically with MaxQuant (v.1.5.5.1 and v.1.6.0.16) using default settings parameters. N-terminal acetylation of protein, methionine oxidation mono-methyl-K/R, di-methyl-K/R and tri-methyl-K were set as variable modifications in MaxQuant. Outputs from MaxQuant were manually filtered as follow: proteins were accepted if identified with at least two peptides of which at least one unique; protein quantified were considered for further analysis only if they have ratio count (RC) equal or greater than 1.

A data analysis pipeline, written in Perl, was developed in-house to process MaxQuant output. In this pipeline, the evidence.txt file was first filtered: potential contaminants and reverse sequences were removed. No Andromeda score or PTM localization probability cut-off was imposed. For quantitative analysis, methyl-peptides ratios were normalised on protein-level H/L information within each IP experiment and the median ratio of redundant methyl-peptides was calculated. To define significantly regulated methyl-peptides upon PRMT1 KD and OE, for each IP experiment, we calculated the mean and standard deviation of the unmodified peptidome distributions. We then calculated the average ratios and standard deviations among biological replicates for each IP experiment and used these parameters to define statistically significant regulated peptides.

Localization analyses of methylated peptides were performed using the Pfam database (http://pfam.xfam.org/), the Eukaryotic Linear Motif (ELM) resource (http://elm.eu.org/) or SMART database (http://smart.embl-heidelberg.de/smart/).

Finally, we employed an in-house developed Perl tool, named hmLINKER, to intersect the SILAC dataset and the previously acquired hmSILAC dataset. For each methyl-peptide identified in the SILAC experiment, hmLINKER checks if a peptide with the same sequence is present in the hmSILAC dataset. In case a match is not found, it then tries to validate the individual modification sites.

### Total and small-RNA extraction

Total and small-RNA were prepared using mirVANA^TM^ miRNA Isolation kit (Ambion), according to the manufacturer’s specification. For the analysis of small-RNA in distinct cellular compartments, RNA was isolated from the chromatin fraction, the nucleosol and the cytosol by using Trizol (Trizol LS Reagent, Ambion). DNAse (Zymo Research) treatment of RNA was performed before reverse transcription. For qPCR analyses of LDC transcriptome, RNA was extracted using the RNA extraction kit (Zymo research) following manufacturer’s instruction.

### cDNA synthesis and qPCR

The cDNA for mRNA, pri-miRNA, pre-miRNA and miRNA profiling was produced using the reverse-transcriptase miScript II RT Kit (Qiagen) according to the manufacturer’s specifications. One tenth of the reaction was used for qPCR reactions in a 7,500HT Fast Real-Time PCR System. miScript SYBR Green PCR Kit (Qiagen) was used for miRNA and pre-miRNA analysis, following manufacturer’s instruction, while mRNA and pri-miRNA were analyzed with FAST Sybr Green Master Mix (Life Technologies).

### Primers used for qPCR

MicroRNAs were amplified using forward primers from miScript primer assays (Qiagen) and reverse universal primers from miScript SYBR Green PCR Kit (Qiagen). For precursor miRNAs analysis, we adapted the miScript Qiagen strategy also to precursor amplification using custom primers. For primary miRNAs, primers were designed to amplify the region upstream the first stem loop of the miRNA cluster.

DGCR8: Fwd 5’ –AAAACTTGCGAAGAATAAAGCTG– 3’,

DGCR8: Rev 5’ –TCTGTTTAACAAAGTCAGGGATGA– 3’

pri-miR-15a/16-1: Fwd 5’ –GCCCTGTTAAGTTGGCATAGC– 3’

pri-miR-15a/16-1: Rev 5’ –ACTGAAGTCCATTCTGTGCCC– 3’

pri-miR-17-92: Fwd 5’ –TGCCACGTGGATGTGAAGAT– 3’

pri-miR-17-92: Rev 5’ –GGCCTCTCCCAAATGGATTGA– 3’

pre-miR-15: Fwd 5’ –CCTTGGAGTAAAGTAGCAGCACA– 3’

pre-miR-16: Fwd 5’ –CAGTGCCTTAGCAGCACGTA– 3’

pre-miR-17: Fwd 5’–AAAGTGCTTACAGTGCAGGTAGT– 3’

pre-miR-18: Fwd 5’ –GGTGCATCTAGTGCAGATAGTGA– 3’

pre-miR-19a: Fwd 5’ –GTCCTCTGTTAGTTTTGCATAGTTG– 3’

pre-miR-19b: Fwd 5’ –GTTAGTTTTGCAGGTTTGCATCC– 3’

pre-miR-20: Fwd 5’ –GTAGCACTAAAGTGCTTATAGTGCAGG– 3’

pre-miR-92: Fwd 5’ –CTACACAGGTTGGGATCGGT– 3’

### Taqman Array Human microRNAs

TaqMan Array Human microRNA A+B Cards (Applied Biosystems) were used for global miRNAs analysis, following manufacturer’ specifications. Data were normalized on the geometric mean of two housekeeping genes (MammU6, U6snRNA). MicroRNAs were considered significantly up/down-regulated when miRNA expression in PRMT1 KD cells relative to the control was greater/lower than 1.5-fold changes, respectively.

### Small-RNA sequencing

Total RNA extracted from PRMT1 KD (sh-1) and control cells was used as input for libraries preparation with the TruSeq Small RNA Library Prep Kit (Illumina), following manufacturer’s instructions. Briefly, RNA 3’ and RNA 5’ adapters were sequentially ligated to the RNA. Reverse transcription followed by PCR was used to create cDNA constructs based on the small RNA ligated with the adapters. The resulting cDNA constructs were gel-purified, eluted and concentrated by ethanol precipitation. The DNA fragment library was quantified with Bioanalyzer (Agilent Technologies) and sequencing was performed on an Illumina HiSeq2000 at 50bp single-read mode and 20 million read depth. Data analysis was performed with the IsomiRage workflow, as previously described (Mueller H.J. et al, 2014). Data normalization was performed after reads mapping, assignment and filtering. Normalization of the data was performed with a reads-per-million (RPM) normalization, using small nucleolar RNAs (snoRNA) reads in each sample as normalizer.

### UV-crosslinked RNA immunoprecipitation (UV-RIP)

UV-RIP protocol was modified from Jeon and Lee, 2001 and Cabianca et al., 2012. Briefly, HeLa cells were harvested and UV-crosslinked with 2 cycles of irradiations at 100000 μJ/cm^2^.Cells were lysed with Lysis Buffer (0.5% NP-40, 0.5% NaDeoxycholate, 1x Roche protease inhibitors mixture (04693116001 MERCK), 25 U/mL Superase-RNAse inhibitor (Thermo Fisher Scientific) in PBS for 30 min at 4°C and then treated with 30 U of Turbo DNAseI (Thermo Fisher Scientific) for 30 min at 37°C. An aliquot (10%) of DNA-digested lysates was used as input while the remaining protein extract (90%) was split in two fractions and incubated overnight at 4°C with either IgG (Millipore) or anti-ILF3 (Bethyl) rabbit antibodies. The day after, protein G dynabeads (Thermo Fisher Scientific) were added and samples rocked for additional 3 hours at 4°C. Afterwards, the dynabeads were washed 4 times with Washing Buffer I (PBS supplemented with 1% NP-40, 0.5% NaDeoxycholate, 300 mM NaCl, 1x Roche protease inhibitor mix and 25 U/mL Superase-RNAse inhibitor), resuspended in RNAse-free water and treated again with Turbo DNAseI for 30 min at 37°C. The input material was treated in parallel in the same manner. The dynabeads were then washed 4 times with Washing Buffer II (PBS supplemented with 1% NP-40, 0.5% NaDeoxycholate, 300 mM NaCl, 10 mM EDTA, 1xRoche protease inhibitor and 25 U/mL Superase-RNAse inhibitor). Finally, the RNA was eluted from the beads with the Elution Buffer (100 mM Tris-HCl pH 7.5, 50 mM NaCl, 10 mM EDTA, 500 μg Proteinase K, 0.5% SDS), for 1 hour at 55 °C. Beads were then pelleted, the supernatants containing the first RNA eluted fraction collected in a clean Eppendorf tube and beads were resuspended in RNA-Lysis Buffer (Zymo Research), to collect additional RNA remained attached to the beads. The two RNA-collected fractions were extracted separately with the RNA-extraction kit (Zymo Research) and subsequently pooled together. Finally, to remove residual genomic DNA, the RNA fractions and the input material were boiled for 2 min at 95°C, followed by 5 min incubation in ice, and then put at 37°C for 1 h in the presence of Turbo DNAseI. The RNA was finally extracted, retro-transcribed to cDNA using the miSCRIPT II RT kit (Qiagen) and analysed by qPCR, as above described. The values obtained for each immunopurifications were normalized over their respective input material and plotted in a histogram, as relative fold enrichment over the not immuno-purified RNA.

### Immunofluorescence

Cells were plated on glass coverslips, fixed in 4% paraformaldehyde for 20 min at room temperature (RT) and permeabilized with PBS-0.1% Triton X-100 for 2 min on ice. Subsequently, cells were initially incubated with PBS-2% BSA for 30 min at RT and later on with the following antibodies dissolved in PBS-2% BSA for 2 hours: anti-EWS (5μg/ml); anti-TAF15 (1:500). After being washed, cells were stained with the respective Alexa Fluor 488 secondary antibody (Molecular Probes, Eugene, OR, USA) diluted 1:400 in PBS-2% BSA for 1 hour at RT. Nuclei were stained with 4′,6-diamidino-2-phenylindole (DAPI). Images were obtained with a Leica TCS SP2 (Leica Microsystems, Heerbrugg, Switzerland).

## Acknowledgments

We thank the Structural Genomic Consortium (SGC, Toronto, Canada) for providing the MS023 and MS094 compounds, and all members of the T. Bonaldi laboratory for helpful discussion.

## Author contributions

V.S. performed the experiments, analyzed the data and wrote the manuscript. R.G performed part of the experiments and the data analysis and wrote the manuscript. F.P. performed some of the wet experiments. M. M. supervised the initial phase of V. S. work. E.M. analyzed the proteomics data. F.G. and F.N. performed the miRNA-seq data analysis. T.B. revised the research project, supervised the experiments, managed the scientific collaboration with F. N. and wrote and edited the manuscript. All authors read and approved the final manuscript.

## Data and Code Availability

MS-proteomics data have been deposited in the ProteomeXchange Consortium via PRIDE [94] partner repository with the dataset identifier PXD011617.

The hmSEEKER code is available on Bitbucket (https://bit.ly/2scCT9u). Perl codes used to analyze the SILAC data and to intersect it with the hmSEEKER output are available upon request.

## References

1. Vaucheret H, Vazquez F, Crete P, Bartel DP (2004) The action of ARGONAUTE1 in the miRNA pathway and its regulation by the miRNA pathway are crucial for plant development. Genes Dev 18: 1187–1197

2. Long JP, Santner TJ, Bartel DL (2009) Hip resurfacing increases bone strains associated with short-term femoral neck fracture. J Orthop Res 27: 1319–1325

3. Ambros V (2004) The functions of animal microRNAs. Nature 431: 350–355

4. Gregory RI, Yan KP, Amuthan G, Chendrimada T, Doratotaj B, Cooch N, Shiekhattar R (2004) The Microprocessor complex mediates the genesis of microRNAs. Nature 432: 235–240

5. Lee YK, Moore DD (2002) Dual mechanisms for repression of the monomeric orphan receptor liver receptor homologous protein-1 by the orphan small heterodimer partner. J Biol Chem 277: 2463–2467

6. Ha M, Kraushaar DC, Zhao K (2014) Genome-wide analysis of H3.3 dissociation reveals high nucleosome turnover at distal regulatory regions of embryonic stem cells. Epigenetics Chromatin 7: 38

7. Denli AM, Tops BB, Plasterk RH, Ketting RF, Hannon GJ (2004) Processing of primary microRNAs by the Microprocessor complex. Nature 432: 231–235

8. Han J, Lee Y, Yeom KH, Kim YK, Jin H, Kim VN (2004) The Drosha-DGCR8 complex in primary microRNA processing. Genes Dev 18: 3016–3027

9. Han J, Lee Y, Yeom KH, Nam JW, Heo I, Rhee JK, Sohn SY, Cho Y, Zhang BT, Kim VN (2006) Molecular basis for the recognition of primary microRNAs by the Drosha-DGCR8 complex. Cell 125: 887–901

10. Blaszczyk J, Tropea JE, Bubunenko M, Routzahn KM, Waugh DS, Court DL, Ji X (2001) Crystallographic and modeling studies of RNase III suggest a mechanism for double-stranded RNA cleavage. Structure 9: 1225–1236

11. Zeng Y, Yi R, Cullen BR (2005) Recognition and cleavage of primary microRNA precursors by the nuclear processing enzyme Drosha. EMBO J 24: 138–148

12. Auyeung VC, Ulitsky I, McGeary SE, Bartel DP (2013) Beyond secondary structure: primary-sequence determinants license pri-miRNA hairpins for processing. Cell 152: 844–858

13. Ma H, Wu Y, Choi JG, Wu H (2013) Lower and upper stem-single-stranded RNA junctions together determine the Drosha cleavage site. Proc Natl Acad Sci U S A 110: 20687–20692

14. Davis BN, Hata A (2009) Regulation of MicroRNA Biogenesis: A miRiad of mechanisms. Cell Commun Signal 7: 18

15. Yi R, Qin Y, Macara IG, Cullen BR (2003) Exportin-5 mediates the nuclear export of pre-microRNAs and short hairpin RNAs. Genes Dev 17: 3011–3016

16. Lund E, Guttinger S, Calado A, Dahlberg JE, Kutay U (2004) Nuclear export of microRNA precursors. Science 303: 95–98

17. Bohnsack MT, Czaplinski K, Gorlich D (2004) Exportin 5 is a RanGTP-dependent dsRNA-binding protein that mediates nuclear export of pre-miRNAs. RNA 10: 185–191

18. Park JE, Heo I, Tian Y, Simanshu DK, Chang H, Jee D, Patel DJ, Kim VN (2011) Dicer recognizes the 5′ end of RNA for efficient and accurate processing. Nature 475: 201–205

19. Chendrimada TP, Gregory RI, Kumaraswamy E, Norman J, Cooch N, Nishikura K, Shiekhattar R (2005) TRBP recruits the Dicer complex to Ago2 for microRNA processing and gene silencing. Nature 436: 740–744

20. Gregory RI, Shiekhattar R (2005) MicroRNA biogenesis and cancer. Cancer Res 65: 3509–3512

21. Li Y, Kowdley KV (2012) MicroRNAs in common human diseases. Genomics Proteomics Bioinformatics 10: 246–253

22. Lin S, Gregory RI (2015) MicroRNA biogenesis pathways in cancer. Nat Rev Cancer 15: 321–333

23. Siomi H, Siomi MC (2010) Posttranscriptional regulation of microRNA biogenesis in animals. Mol Cell 38: 323–332

24. Han J, Pedersen JS, Kwon SC, Belair CD, Kim YK, Yeom KH, Yang WY, Haussler D, Blelloch R, Kim VN (2009) Posttranscriptional crossregulation between Drosha and DGCR8. Cell 136: 75–84

25. Shiohama A, Sasaki T, Noda S, Minoshima S, Shimizu N (2007) Nucleolar localization of DGCR8 and identification of eleven DGCR8-associated proteins. Exp Cell Res 313: 4196–4207

26. Newman MA, Hammond SM (2010) Emerging paradigms of regulated microRNA processing. Genes Dev 24: 1086–1092

27. Fukuda T, Yamagata K, Fujiyama S, Matsumoto T, Koshida I, Yoshimura K, Mihara M, Naitou M, Endoh H, Nakamura T, et al. (2007) DEAD-box RNA helicase subunits of the Drosha complex are required for processing of rRNA and a subset of microRNAs. Nat Cell Biol 9: 604–611

28. Guil S, Caceres JF (2007) The multifunctional RNA-binding protein hnRNP A1 is required for processing of miR-18a. Nat Struct Mol Biol 14: 591–596

29. Kawahara Y, Mieda-Sato A (2012) TDP-43 promotes microRNA biogenesis as a component of the Drosha and Dicer complexes. Proc Natl Acad Sci U S A 109: 3347–3352

30. Di Carlo V, Grossi E, Laneve P, Morlando M, Dini Modigliani S, Ballarino M, Bozzoni I, Caffarelli E (2013) TDP-43 regulates the microprocessor complex activity during in vitro neuronal differentiation. Mol Neurobiol 48: 952–963

31. Sakamoto S, Aoki K, Higuchi T, Todaka H, Morisawa K, Tamaki N, Hatano E, Fukushima A, Taniguchi T, Agata Y (2009) The NF90-NF45 complex functions as a negative regulator in the microRNA processing pathway. Mol Cell Biol 29: 3754–3769

32. Higuchi T, Todaka H, Sugiyama Y, Ono M, Tamaki N, Hatano E, Takezaki Y, Hanazaki K, Miwa T, Lai S, et al. (2016) Suppression of MicroRNA-7 (miR-7) Biogenesis by Nuclear Factor 90-Nuclear Factor 45 Complex (NF90-NF45) Controls Cell Proliferation in Hepatocellular Carcinoma. J Biol Chem 291: 21074–21084

33. Nussbacher JK, Yeo GW (2018) Systematic Discovery of RNA Binding Proteins that Regulate MicroRNA Levels. Mol Cell 69: 1005–1016e1007

34. Herbert KM, Pimienta G, DeGregorio SJ, Alexandrov A, Steitz JA (2013) Phosphorylation of DGCR8 increases its intracellular stability and induces a progrowth miRNA profile. Cell Rep 5: 1070–1081

35. Tang X, Wen S, Zheng D, Tucker L, Cao L, Pantazatos D, Moss SF, Ramratnam B (2013) Acetylation of drosha on the N-terminus inhibits its degradation by ubiquitination. PLoS One 8: e72503

36. Yang Q, Li W, She H, Dou J, Duong DM, Du Y, Yang SH, Seyfried NT, Fu H, Gao G, et al. (2015) Stress induces p38 MAPK-mediated phosphorylation and inhibition of Drosha-dependent cell survival. Mol Cell 57: 721–734

37. Bremang M, Cuomo A, Agresta AM, Stugiewicz M, Spadotto V, Bonaldi T (2013) Mass spectrometry-based identification and characterisation of lysine and arginine methylation in the human proteome. Mol Biosyst 9: 2231–2247

38. Larsen SC, Sylvestersen KB, Mund A, Lyon D, Mullari M, Madsen MV, Daniel JA, Jensen LJ, Nielsen ML (2016) Proteome-wide analysis of arginine monomethylation reveals widespread occurrence in human cells. Sci Signal 9: rs9

39. Yang Y, Bedford MT (2013) Protein arginine methyltransferases and cancer. Nat Rev Cancer 13: 37–50

40. Bedford MT, Clarke SG (2009) Protein arginine methylation in mammals: who, what, and why. Mol Cell 33: 1–13

41. Tang J, Frankel A, Cook RJ, Kim S, Paik WK, Williams KR, Clarke S, Herschman HR (2000) PRMT1 is the predominant type I protein arginine methyltransferase in mammalian cells. J Biol Chem 275: 7723–7730

42. Yu Z, Chen T, Hebert J, Li E, Richard S (2009) A mouse PRMT1 null allele defines an essential role for arginine methylation in genome maintenance and cell proliferation. Mol Cell Biol 29: 2982–2996

43. Bedford MT, Richard S (2005) Arginine methylation an emerging regulator of protein function. Mol Cell 18: 263–272

44. Wada K, Inoue K, Hagiwara M (2002) Identification of methylated proteins by protein arginine N-methyltransferase 1, PRMT1, with a new expression cloning strategy. Biochim Biophys Acta 1591: 1–10

45. Strahl BD, Briggs SD, Brame CJ, Caldwell JA, Koh SS, Ma H, Cook RG, Shabanowitz J, Hunt DF, Stallcup MR, et al. (2001) Methylation of histone H4 at arginine 3 occurs in vivo and is mediated by the nuclear receptor coactivator PRMT1. Curr Biol 11: 996–1000

46. Wang H, Huang ZQ, Xia L, Feng Q, Erdjument-Bromage H, Strahl BD, Briggs SD, Allis CD, Wong J, Tempst P, et al. (2001) Methylation of histone H4 at arginine 3 facilitating transcriptional activation by nuclear hormone receptor. Science 293: 853–857

47. Zhao X, Jankovic V, Gural A, Huang G, Pardanani A, Menendez S, Zhang J, Dunne R, Xiao A, Erdjument-Bromage H, et al. (2008) Methylation of RUNX1 by PRMT1 abrogates SIN3A binding and potentiates its transcriptional activity. Genes Dev 22: 640–653

48. Kwak YT, Guo J, Prajapati S, Park KJ, Surabhi RM, Miller B, Gehrig P, Gaynor RB (2003) Methylation of SPT5 regulates its interaction with RNA polymerase II and transcriptional elongation properties. Mol Cell 11: 1055–1066

49. Boisvert FM, Rhie A, Richard S, Doherty AJ (2005) The GAR motif of 53BP1 is arginine methylated by PRMT1 and is necessary for 53BP1 DNA binding activity. Cell Cycle 4: 1834–1841

50. Boisvert FM, Dery U, Masson JY, Richard S (2005) Arginine methylation of MRE11 by PRMT1 is required for DNA damage checkpoint control. Genes Dev 19: 671–676

51. Dery U, Coulombe Y, Rodrigue A, Stasiak A, Richard S, Masson JY (2008) A glycine-arginine domain in control of the human MRE11 DNA repair protein. Mol Cell Biol 28: 3058–3069

52. Guendel I, Carpio L, Pedati C, Schwartz A, Teal C, Kashanchi F, Kehn-Hall K (2010) Methylation of the tumor suppressor protein, BRCA1, influences its transcriptional cofactor function. PLoS One 5: e11379

53. Belyanskaya LL, Gehrig PM, Gehring H (2001) Exposure on cell surface and extensive arginine methylation of ewing sarcoma (EWS) protein. J Biol Chem 276: 18681–18687

54. Araya N, Hiraga H, Kako K, Arao Y, Kato S, Fukamizu A (2005) Transcriptional down-regulation through nuclear exclusion of EWS methylated by PRMT1. Biochem Biophys Res Commun 329: 653–660

55. Tradewell ML, Yu Z, Tibshirani M, Boulanger MC, Durham HD, Richard S (2012) Arginine methylation by PRMT1 regulates nuclear-cytoplasmic localization and toxicity of FUS/TLS harbouring ALS-linked mutations. Hum Mol Genet 21: 136–149

56. Scaramuzzino C, Monaghan J, Milioto C, Lanson NA, Jr., Maltare A, Aggarwal T, Casci I, Fackelmayer FO, Pennuto M, Pandey UB (2013) Protein arginine methyltransferase 1 and 8 interact with FUS to modify its sub-cellular distribution and toxicity in vitro and in vivo. PLoS One 8: e61576

57. Dammer EB, Fallini C, Gozal YM, Duong DM, Rossoll W, Xu P, Lah JJ, Levey AI, Peng J, Bassell GJ, et al. (2012) Coaggregation of RNA-binding proteins in a model of TDP-43 proteinopathy with selective RGG motif methylation and a role for RRM1 ubiquitination. PLoS One 7: e38658

58. Destouches D, El Khoury D, Hamma-Kourbali Y, Krust B, Albanese P, Katsoris P, Guichard G, Briand JP, Courty J, Hovanessian AG (2008) Suppression of tumor growth and angiogenesis by a specific antagonist of the cell-surface expressed nucleolin. PLoS One 3: e2518

59. Blackwell E, Ceman S (2012) Arginine methylation of RNA-binding proteins regulates cell function and differentiation. Mol Reprod Dev 79: 163–175

60. Stetler A, Winograd C, Sayegh J, Cheever A, Patton E, Zhang X, Clarke S, Ceman S (2006) Identification and characterization of the methyl arginines in the fragile X mental retardation protein Fmrp. Hum Mol Genet 15: 87–96

61. Ong SE, Mittler G, Mann M (2004) Identifying and quantifying in vivo methylation sites by heavy methyl SILAC. Nat Methods 1: 119–126

62. Cumberworth A, Lamour G, Babu MM, Gsponer J (2013) Promiscuity as a functional trait: intrinsically disordered regions as central players of interactomes. Biochem J 454: 361–369

63. Kato M, Han TW, Xie S, Shi K, Du X, Wu LC, Mirzaei H, Goldsmith EJ, Longgood J, Pei J, et al. (2012) Cell-free formation of RNA granules: low complexity sequence domains form dynamic fibers within hydrogels. Cell 149: 753–767

64. Calabretta S, Richard S (2015) Emerging Roles of Disordered Sequences in RNA-Binding Proteins. Trends Biochem Sci 40: 662–672

65. Pekarsky Y, Croce CM (2015) Role of miR-15/16 in CLL. Cell Death Differ 22: 6–11

66. Olive V, Sabio E, Bennett MJ, De Jong CS, Biton A, McGann JC, Greaney SK, Sodir NM, Zhou AY, Balakrishnan A, et al. (2013) A component of the mir-17-92 polycistronic oncomir promotes oncogene-dependent apoptosis. Elife 2: e00822

67. Wang D, Huang J, Hu Z (2012) RNA helicase DDX5 regulates microRNA expression and contributes to cytoskeletal reorganization in basal breast cancer cells. Mol Cell Proteomics 11: M111011932

68. Allegra D, Bilan V, Garding A, Dohner H, Stilgenbauer S, Kuchenbauer F, Mertens D, Zucknick M (2014) Defective DROSHA processing contributes to downregulation of MiR-15/-16 in chronic lymphocytic leukemia. Leukemia 28: 98–107

69. Pickering BF, Yu D, Van Dyke MW (2011) Nucleolin protein interacts with microprocessor complex to affect biogenesis of microRNAs 15a and 16. J Biol Chem 286: 44095–44103

70. Lin SL, Miller JD, Ying SY (2006) Intronic microRNA (miRNA). J Biomed Biotechnol 2006: 26818

71. Lin SL, Chang D, Wu DY, Ying SY (2003) A novel RNA splicing-mediated gene silencing mechanism potential for genome evolution. Biochem Biophys Res Commun 310: 754–760

72. Dhar S, Vemulapalli V, Patananan AN, Huang GL, Di Lorenzo A, Richard S, Comb MJ, Guo A, Clarke SG, Bedford MT (2013) Loss of the major Type I arginine methyltransferase PRMT1 causes substrate scavenging by other PRMTs. Sci Rep 3: 1311

73. Thandapani P, O’Connor TR, Bailey TL, Richard S (2013) Defining the RGG/RG motif. Mol Cell 50: 613–623

74. Eram MS, Shen Y, Szewczyk M, Wu H, Senisterra G, Li F, Butler KV, Kaniskan HU, Speed BA, Dela Sena C, et al. (2016) A Potent, Selective, and Cell-Active Inhibitor of Human Type I Protein Arginine Methyltransferases. ACS Chem Biol 11: 772–781

75. Ozdilek BA, Thompson VF, Ahmed NS, White CI, Batey RT, Schwartz JC (2017) Intrinsically disordered RGG/RG domains mediate degenerate specificity in RNA binding. Nucleic Acids Res 45: 7984–7996

76. Geoghegan V, Guo A, Trudgian D, Thomas B, Acuto O (2015) Comprehensive identification of arginine methylation in primary T cells reveals regulatory roles in cell signalling. Nat Commun 6: 6758

77. Hart-Smith G, Yagoub D, Tay AP, Pickford R, Wilkins MR (2016) Large Scale Mass Spectrometry-based Identifications of Enzyme-mediated Protein Methylation Are Subject to High False Discovery Rates. Mol Cell Proteomics 15: 989–1006

78. Sylvestersen KB, Horn H, Jungmichel S, Jensen LJ, Nielsen ML (2014) Proteomic analysis of arginine methylation sites in human cells reveals dynamic regulation during transcriptional arrest. Mol Cell Proteomics 13: 2072–2088

79. Guo A, Gu H, Zhou J, Mulhern D, Wang Y, Lee KA, Yang V, Aguiar M, Kornhauser J, Jia X, et al. (2014) Immunoaffinity enrichment and mass spectrometry analysis of protein methylation. Mol Cell Proteomics 13: 372–387

80. Hung CJ, Lee YJ, Chen DH, Li C (2009) Proteomic analysis of methylarginine-containing proteins in H.L. cells by two-dimensional gel electrophoresis and immunoblotting with a methylarginine-specific antibody. Protein J 28: 139–147

81. Jobert L, Argentini M, Tora L (2009) PRMT1 mediated methylation of TAF15 is required for its positive gene regulatory function. Exp Cell Res 315: 1273–1286

82. Rajpurohit R, Paik WK, Kim S (1994) Effect of enzymic methylation of heterogeneous ribonucleoprotein particle A1 on its nucleic-acid binding and controlled proteolysis. Biochem J 304 (Pt 3): 903–909

83. Chong PA, Vernon RM, Forman-Kay JD (2018) RGG/RG Motif Regions in RNA Binding and Phase Separation. J Mol Biol 430: 4650–4665

84. Baldwin RM, Morettin A, Paris G, Goulet I, Cote J (2012) Alternatively spliced protein arginine methyltransferase 1 isoform PRMT1v2 promotes the survival and invasiveness of breast cancer cells. Cell Cycle 11: 4597–4612

85. Yoshimatsu M, Toyokawa G, Hayami S, Unoki M, Tsunoda T, Field HI, Kelly JD, Neal DE, Maehara Y, Ponder BA, et al. (2011) Dysregulation of PRMT1 and PRMT6, Type I arginine methyltransferases, is involved in various types of human cancers. Int J Cancer 128: 562–573

86. Bao J, Di Lorenzo A, Lin K, Lu Y, Zhong Y, Sebastian MM, Muller WJ, Yang Y, Bedford MT (2018) Mouse models of overexpression reveal distinct oncogenic roles for different type I protein arginine methyltransferases. Cancer Res, 10.1158/0008-5472.CAN-18-1995

87. Lu J, Getz G, Miska EA, Alvarez-Saavedra E, Lamb J, Peck D, Sweet-Cordero A, Ebert BL, Mak RH, Ferrando AA, et al. (2005) MicroRNA expression profiles classify human cancers. Nature 435: 834–838

88. Calin GA, Ferracin M, Cimmino A, Di Leva G, Shimizu M, Wojcik SE, Iorio MV, Visone R, Sever NI, Fabbri M, et al. (2005) A MicroRNA signature associated with prognosis and progression in chronic lymphocytic leukemia. N Engl J Med 353: 1793–1801

89. Thomson JM, Newman M, Parker JS, Morin-Kensicki EM, Wright T, Hammond SM (2006) Extensive post-transcriptional regulation of microRNAs and its implications for cancer. Genes Dev 20: 2202–2207

90. Shevchenko A, Tomas H, Havlis J, Olsen JV, Mann M (2006) In-gel digestion for mass spectrometric characterization of proteins and proteomes. Nat Protoc 1: 2856–2860

91. Rappsilber J, Mann M, Ishihama Y (2007) Protocol for micro-purification, enrichment, prefractionation and storage of peptides for proteomics using StageTips. Nat Protoc 2: 1896–1906

92. Cox J, Mann M (2008) MaxQuant enables high peptide identification rates, individualized p.p.b.-range mass accuracies and proteome-wide protein quantification. Nat Biotechnol 26: 1367–1372

93. Cox J, Neuhauser N, Michalski A, Scheltema RA, Olsen JV, Mann M (2011) Andromeda: a peptide search engine integrated into the MaxQuant environment. J Proteome Res 10: 1794–1805

94. Vizcaino JA, Csordas A, Del-Toro N, Dianes JA, Griss J, Lavidas I, Mayer G, Perez-Riverol Y, Reisinger F, Ternent T, et al. (2016) 2016 update of the PRIDE database and its related tools. Nucleic Acids Res 44: 11033

95. Letunic I, Bork P (2018) 20 years of the SMART protein domain annotation resource. Nucleic Acids Res 46: D493–D496

96. Hinske LC, Franca GS, Torres HA, Ohara DT, Lopes-Ramos CM, Heyn J, Reis LF, Ohno-Machado L, Kreth S, Galante PA (2014) miRIAD-integrating microRNA inter- and intragenic data. Database (Oxford) 2014:

97. Chou MF, Schwartz D (2011) Biological sequence motif discovery using motif-x. Curr Protoc Bioinformatics Chapter 13: Unit 13 15–24

